# How cBAF and PBAF regulate the nucleosomal and subnucleosomal landscape of promoters, enhancers and REST repressor binding sites

**DOI:** 10.1101/2025.06.12.659349

**Authors:** Manon Soleil, Emilie Drouineau, Marina C. Nocente, Anida Mesihovic Karamitsos, Yuliia Kovalchuk, Hélène Picaud, Cécile Dulary, Sophie Chantalat, Matthieu Gérard

**Affiliations:** Université Paris-Saclay, CEA, CNRS, Institute for Integrative Biology of the Cell (I2BC), 91198 Gif-sur-Yvette, France; Institut de biologie François Jacob, Centre National de Recherche en Génomique Humaine, 91057 Evry, France

## Abstract

BAF (SWI/SNF) chromatin remodeling complexes belong to three subfamilies, cBAF, PBAF, and ncBAF, that regulate the nucleosomal organization and accessibility of transcription factors (TFs) DNA binding sites at cis-regulatory elements (CREs). However, the functional specificities of these complexes at promoters, enhancers, and other categories of CREs are poorly understood. Here, we examined how the BAF complexes regulate the nucleosomal and subnucleosomal landscape in which transcription initiation occurs in vivo. We first described how these complexes regulate the earliest step of transcription preinitiation complex (PIC) assembly, which is defined by the binding of the general transcription factors TBP and TFIIA. We show that in the presence of a TATA box, PIC formation occurs onto a core promoter or enhancer region still wrapped into a fragile nucleosome or a subnucleosomal particle. Conversely, without a TATA box, PIC assembles onto genomic templates mostly stripped of histones. We establish that cBAF, but not PBAF or ncBAF, regulates this chromatin organization and the first step of PIC formation at promoters and enhancers. We observed that individual subunits of cBAF have specific functions at promoters: the BRG1 ATPase and SMARCB1 regulate the production of fragile nucleosomes and subnucleosomal particles, while ARID1A stimulates the binding of TBP/TFIIA independently of the ATPase subunit. In contrast, all three subunits are systematically required for both chromatin organization and TBP/TFIIA binding at enhancers. Finally, we revealed that a subset of PBAF subunits controls the nucleosomal and subnucleosomal landscape at REST repressor binding sites. Our integrative study identified specific functions of cBAF and PBAF subunits in controlling the binding of TFs and the nucleosomal and subnucleosomal organization of CREs.

## INTRODUCTION

The transcription of mammalian genes requires the assembly of the preinitiation complex (PIC) at core promoters. The PIC includes the TFIID, TFIIA, TFIIB, TFIIF, TFIIE, and TFIIH general transcription factors (GTFs) and Pol II, which assemble in a stepwise manner ^1^. TFIID is a large multiprotein complex of ∼1.5 MDa, containing the TATA-box binding protein (TBP) associated with 13 other TBP-associated factors (TAFs)^2^. The core promoter is a region extending 50 bp up- and downstream of the transcription start site (TSS) of each gene. This interval contain a variable combination of DNA motifs including the TATA box (or TBP binding motif), the initiator (INR), the TCT initiator, the downstream promoter element (DPE), Motif Ten Element (MTE), and TFIIB recognition elements^3–6^.

Two main classes of promoters were identified based on the pattern of transcription initiation^7^. The first class is composed of sharp or focused promoters, which have a well-defined TSS position restricted to a few base pairs. The second class is characterized by a broad or dispersed TSS position, in which a series of TSS is typically distributed within a ∼100-bp window. Sharp promoters are more likely to contain a TATA box and other known core promoter elements than broad promoters^8^. The TATA box is however present only in ∼10% of the sharp promoters, and is thus not the sole determinant of the focused mode of transcription initiation. Broad promoters are often associated with CpG-island and are relatively depleted of the consensus DNA motifs described above compared to sharp promoters.

Structural analysis suggest that PIC assembly is initiated by the binding of TFIID to the core promoter^9^. During this early step of PIC formation, TFIID experiences drastic conformational rearrangements resulting in the proper positioning of TBP and TFIIA on the promoter^10–12^. Core promoters containing both a TATA box and a DPE provide two anchoring point that stabilize TFIID binding during PIC formation and transcription initiation. This stabilization of TFIID was proposed to be the key mechanism controlling a focused transcription initiation^3^. Promoters lacking a TATA box or a DPE would allow a more labile interaction of TFIID, increasing the probability of TFIID repositioning relative to the core promoters, resulting in the broad initiation pattern^3^.

Most of the studies on the formation of the PIC have been conducted using DNA templates devoid of nucleosomes. In the nucleus, promoters have a well-defined nucleosomal organization, in which two positioned nucleosomes flank a nucleosome free region (NFR), also called a nucleosome depleted region (NDR)^13–15^. The NFR spans a few tens to several hundreds of bp and includes the TSS^15^, which is thus in principle accessible to the GTFs and Pol II to initiate PIC formation. However, we and others have detected “fragile”, MNase-accessible nucleosomes within the NFR ^16–21^. The TSS is often embedded within the fragile nucleosome^21^, suggesting that the accessibility of the TSS is regulated by these particles. The DNA region wrapped into the fragile nucleosome presumably experiences frequent remodeling events, as partially unwrapped nucleosomes and subnucleosomal particles were detected at the same genomic position^21,22^.

The nucleosomal and subnucleosomal organization, as well as the formation of the NFR are influenced by the activities of a large class of enzymes called ATP-dependent chromatin remodelers ^23,24^. We and others have shown that the BAF (BRG1/BRM-Associated Factors, or SWI/SNF) complexes are critically involved in controlling the chromatin landscape of enhancers^7,21,22,25–27^. BAF complexes regulate the binding of a variety of TFs that are not all binding to enhancer elements, including the REST (Repressor element 1 silencing transcription factor) transcriptional repressor^28^. REST is a zinc finger-containing TF that binds to its specific DNA binding motif to silence neuronal genes in non-neuronal cells^29^.

The BAF complexes consist of three distinct families, defined by their protein subunit composition: canonical BAF (cBAF, 12 subunits), polybromo-associated BAF (PBAF, 13 subunits), and non-canonical BAF (ncBAF, 10 subunits)^30,31^. Eight subunits are shared among the three complexes, forming the initial assembly module and the ATPase module, the latter containing one of the two mutually exclusive ATPases, BRG1 or BRM. Some subunits, like SMARCB1, are shared between cBAF and PBAF, while others are unique to each complex. cBAF contains one of two AT-rich interaction domain (ARID) subunits, ARID1A or ARID1B, and a DPF family protein (DPF1, 2, or 3). PBAF includes ARID2 and two bromodomain proteins, PBRM1 and BRD7. Finally, ncBAF incorporates two specific subunits, BRD9 and GLTSCR1 or GLTSCR1L (reviewed in reference^30^).

In this study, we have explored in detail the nucleosomal and subnucleosomal organization of the core promoter region in mouse embryonic stem (ES) cells, and defined how components of the PIC interact with this *in vivo* environment. We observed that early PIC formation occurs at the genomic location in which we detect the fragile nucleosomes and subnucleosomal particles. We further demonstrated that these fragile nucleosomes and subnucleosomes co-occur with the binding of the GTFs of the early PIC at TATA-box containing promoters. We have analyzed how the BAF complexes regulate the early phase of PIC formation at core promoters and enhancers. We observed that the cBAF complex is specifically involved in regulating PIC formation at these cis-regulatory elements (CRE). However, the importance of cBAF is different at enhancers and promoters. While most subunits of cBAF are critically required for PIC formation at enhancers, cBAF has only a stimulatory function on TBP and TFIIA binding to the core promoter. This distinct regulation of PIC formation correlates with a differential control by cBAF of the nucleosomal and subnucleosomal organization at enhancers and promoters. Our investigations failed to identify a function for PBAF and ncBAF in regulating PIC formation. However, we detected a highly specific function of PBAF in regulating the nucleosomal and subnucleosomal landscape of REST repressor binding sites.

## RESULTS

### Nucleosomal and subnucleosomal landscape of transcription initiation in ES cells

We first compared the nucleosomal and subnucleosomal organization of broad and sharp promoters by MNase ChIP-seq, using antibodies against histone H3. We found that both categories of promoters have a similar organization, in which two canonical nucleosomes labeled -2 and +1 relative to the TSS flank the NFR (Fig. 1). Importantly, the NFR contains a nucleosome hypersensitive to MNase digestion (thus called a fragile nucleosome) at position -1 (reference^21^). The position of the dyad axis of the fragile nucleosome is located immediately upstream of the TSS (Fig. 1a). Because of their adjacent positions, the TSS is often embedded within the fragile nucleosome, suggesting a reduced accessibility of the TSS in this context. We detected a TATA box in about 9% of the sharp and 3% of the broad promoters (Methods). The presence of a TATA box did not modify the nucleosomal organization of the promoter, compared to broad promoters or TATA-less sharp promoters (Fig. 1a).

**Figure 1.**
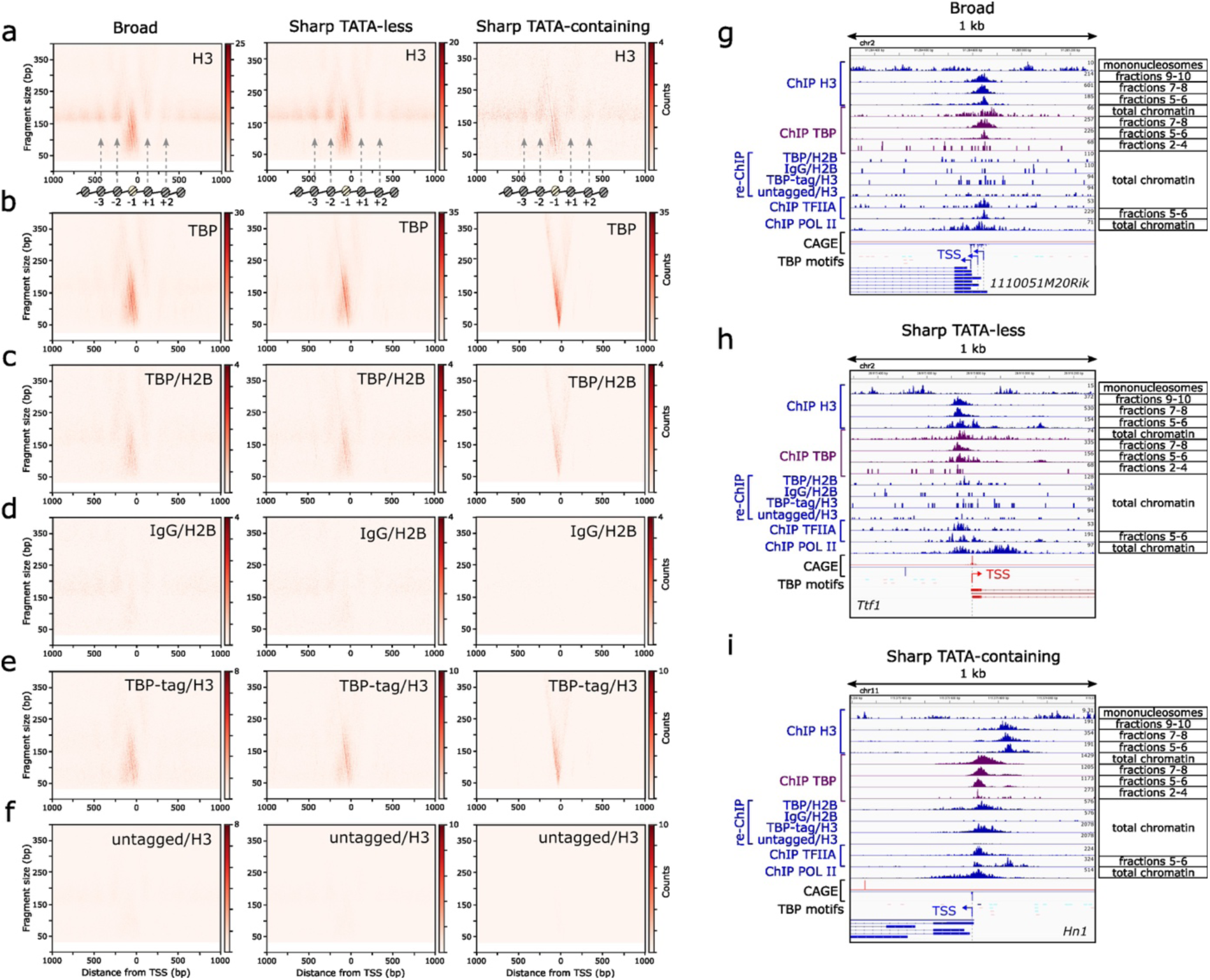
Nucleosomal and subnucleosomal landscape of transcription initiation in ES cells. **a, b**. V-plots of histone H3 (**a**) and TBP (**b**) ChIP-seq fragments spanning ±1,000 bp from the Transcription Start Site (TSS). Red dots indicate the genomic position of the midpoint of each immunoprecipitated DNA fragment on the x axis and its length in bp on the y axis. The color scale corresponds to the number of DNA fragments. The schematic illustration in (a) indicates the positions of the canonical nucleosomes (grey). The -2 and +1 nucleosomes flank either the fragile nucleosome - 1 (yellow) or the subnucleosomal particles that occupy the central region. **c-f**. Sequential ChIP-seq experiments. **c, d**. Chromatin from WT mouse ES cells was first immunoprecipitated with a TBP monoclonal antibody (**c**) or an unspecific IgG (**d**). Elution was performed by peptide competition and each eluted fraction was subjected to a second round of immunoprecipitation with an antibody against H2B. **e, f**. Chromatin from TBP-tagged (**e**) or untagged mouse ES cells (**f**) was first immunoprecipitated with an HA antibody. Elution was performed by TEV cleavage and each eluted fraction was subjected to a second round of immunoprecipitation with an antibody against H3. For panels (**a-f**), the promoters are separated according to their shape (broad or sharp) and the presence or absence of the TATA box. **g-i**. Density graphs showing the distribution of ChIP DNA fragment centers at representative examples of the three types of promoters. The top lane shows the positions of canonical nucleosomes detected by histone H3 ChIP-seq of sucrose gradient fractions 11 and 13. The three following lanes show the position of the fragile nucleosomes (fractions 9-10) and the different subnucleosomal species (fractions 7-8 and 5-6), revealed by histone H3 ChIP-seq of sucrose gradient fractions (see figure 2). The following lanes show TBP ChIP-seq signal of total chromatin, subnucleosomal particles (pools of sucrose gradient fractions 7-8 and 5-6), and histone-free DNA (fractions 2-4). The four subsequent lanes represent sequential ChIP-seq signal corresponding respectively to (**c**), (**d**), (**e**) and (**f**). The following lanes show TFIIA ChIP-seq signals of total chromatin and fractions 5-6. The subsequent lane represents Pol II ChIP-seq signal on total chromatin. CAGE signal maps the position of the TSS. TBP consensus motifs with high (P = 0.001) and low (P = 0.01) confidence are indicated, the darker the color, the higher the confidence. For CAGE, TATA motifs and annotations, the color corresponds to the gene orientation: the genes depicted in blue are oriented to the left, while the genes depicted in red are oriented to the right. The TSS is symbolized by a unique arrow for sharp promoters and multiple arrows for broad promoters. At least two biological replicates were performed for each ChIP-seq experiment.

We previously identified a variety of subnucleosomal particles sharing the genomic position of the fragile nucleosome at enhancers and promoters, using a combination of MNase ChIP-seq and sucrose gradient centrifugation (referred to as gradient ChIP-seq hereafter)^21^. We detected a similar subnucleosomal organization at sharp and broad promoters: the fragile nucleosome had a slightly lower sedimentation rate than the canonical nucleosomes flanking the NFR and was detected in fractions 9-10 of the gradient (Fig. 2a, b). As expected from our previous study, we observed subnucleosomal particles of smaller apparent molecular weight in fractions 5-6 and 7-8 (Fig. 2c, e). After the centrifugation step, the top of the gradient fractions (fractions 2-4) contained mostly histone-free DNA fragments protected from MNase digestion by formaldehyde-crosslinked TF (reference^21^ and Fig. 2g). In conclusion, broad and sharp promoters share a similar subnucleosomal organization, irrespective of the presence of a TATA box.

**Figure 2.**
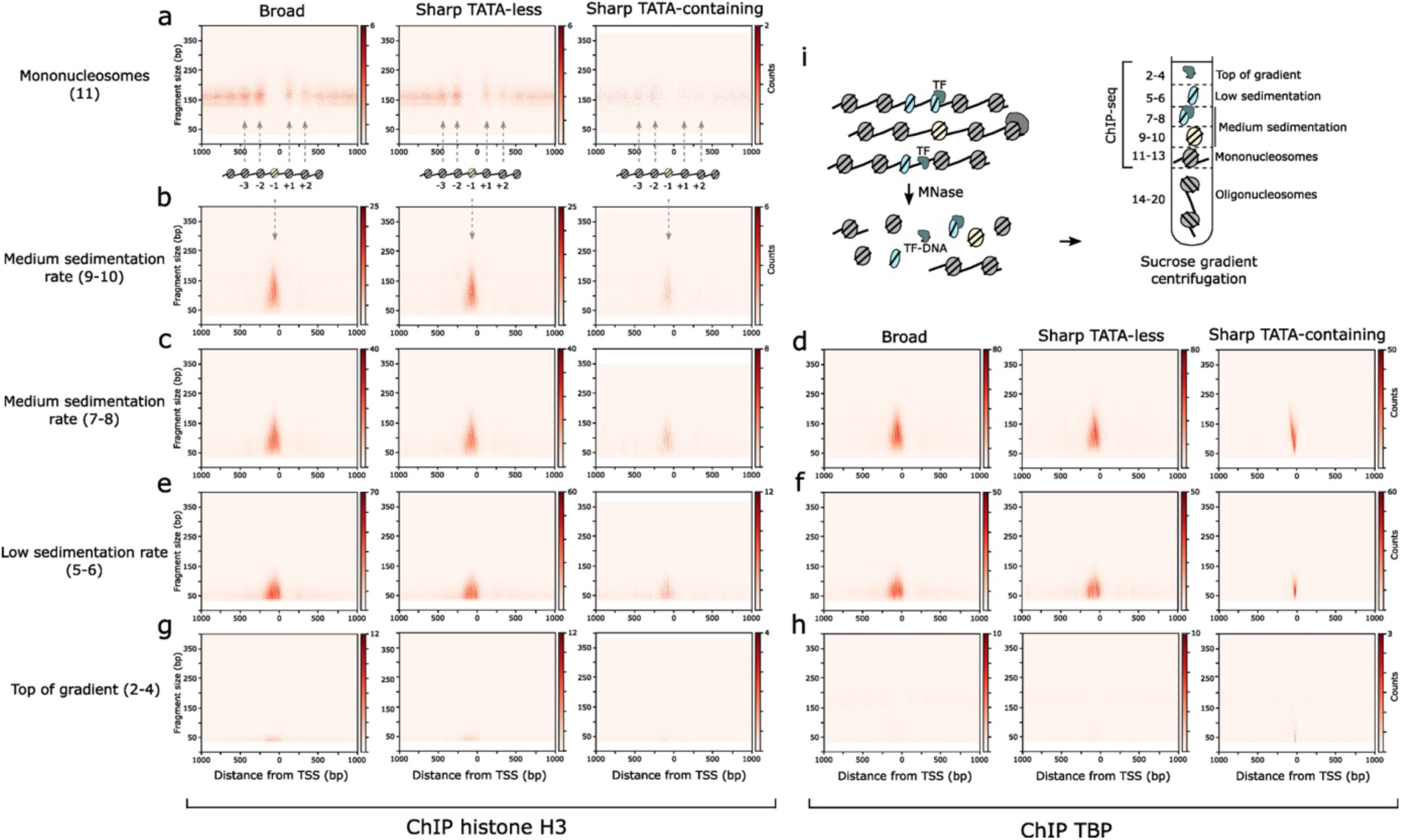
TBP binds the subnucleosomal particles of the core promoter. The chromatin from formaldehyde-fixed ES cells was digested with a moderate dose of MNase and centrifugated through a 10–30% sucrose gradient to separate canonical nucleosomes from subnucleosomal particles and TF–DNA complexes (**i**). **a-h**. V-plots of histone H3 (**a-c, e, g**) and TBP (**d, f, h**) ChIP-seq experiments performed with sucrose gradient fractions as the input, spanning ±1,000 bp from the TSS. The fractions or pools used for ChIP are indicated on the left, and the type of promoter on the top. The schematic illustration indicates the positions of the canonical nucleosomes (**a**) and the fragile nucleosome -1 (**b**), as in Figure 1. At least two biological replicates were performed for each ChIP-seq experiment.

### TBP binds the TATA box in the context of the fragile nucleosome and its associated subnucleosomal particle

Next, we investigated how two components of the early PIC, TBP and TFIIA, integrate the nucleosomal and subnucleosomal landscape of the promoters. We observed that TBP binding occurs in the same region of the core promoter in which we detect the fragile nucleosomes and subnucleosomal particles (Fig. 1b). The TBP binding pattern was however more dispersed in the broad than in the sharp promoter category, and was the most focused in the TATA-containing promoter subclass (Fig. 1b).

We wondered whether TBP could interact with the core promoter DNA template engaged in interactions with histones. To test this possibility, we performed sequential ChIP experiments using a monoclonal antibody against TBP in the first ChIP step, elution by peptide competition, and a second (re)ChIP step using an antibody against histone H2B. This experiment revealed that at least a fraction of TBP interacts with the upstream core promoter region associated with histones (Fig. 1c, d). We confirmed the result of this experiment with an ES cell line expressing an HA-tagged version of TBP. Using this cell line as a source of chromatin, we performed a second series of sequential ChIP experiments with a monoclonal antibody against the HA epitope in the first step, followed by elution by tobacco etch virus (TEV) protease cleavage and a reChIP step using an antibody against histone H3 (Fig. 1e, f). This last experiment confirmed that in vivo, a fraction of TBP binds the core promoter wrapped within a fragile nucleosome or a subnucleosomal particle.

Next, we examined individual genes to appreciate potential variations of this binding pattern across the different classes of promoters. This investigation revealed that 86% of the TATA-box containing, but only 22% of the TATA-less promoters exhibit a high reChIP TBP/histone signal (Fig. 1g-i and Extended Data Fig. 1). Thus, binding of TBP onto TATA-containing promoters in more likely to occur in the context of a chromatinized core promoter than at TATA-less promoters. We investigated whether these chromatinized templates might correspond to the subnucleosomal particles we detected at the core promoter. We used the gradient ChIP-seq method to test the association of TBP with medium or low sedimentation rate subnucleosomal particles and with histone-free DNA (Fig. 2d-h). For all categories of promoters, we detected TBP associated with both the low and medium sedimentation particles, suggesting that TBP can interact in vivo with the subnucleosomes generated at the core promoter (Fig. 2d, f). TBP association with histone-free DNA was only detected at a subset of TATA-containing promoters (Fig. 2h, Fig. 1i, and Extended Data Fig. 1). The binding of TBP was often, but not systematically, correlated with the binding of TFIIA. TFIIA could also be detected bound to low sedimentation particles (Fig. 1g-i and Extended Data Fig. 1). At TATA-containing promoters, TBP and TFIIA binding occurred at the position of the TATA box, suggesting that we had captured by ChIP-seq the first step of PIC formation (Fig. 1g-i and Extended Data Fig. 1).

Structural data suggest that the formation of the PIC is initiated by the binding of TFIID to the core promoter, through interactions with the TATA box and the DPE when both elements are available, or the DPE alone at TATA-less promoters. During this early step of PIC formation, TFIID experiences dramatic conformational rearrangements leading to the proper positioning of TBP and TFIIA on the promoter^10–12^. At the end of these rearrangements, TBP remains connected to the other subunits of TFIID solely through its interaction with TFIIA. It is unclear whether the rest of the TFIID subunits remain associated with the core promoter after this stage. Our ChIP-seq gradient data revealed that TBP is detected within genomic particles having low and medium sedimentation rates at both sharp and broad promoters, irrespective of the presence of a TATA box. Low and medium sedimentation particles have apparent molecular masses of ∼100 and <200 kDa, respectively, well below the ∼1.3 MDa molecular mass of TFIID. These data, which indicate that at least a proportion of TBP is present on the core promoter independently of the other subunits of TFIID, suggest a scenario in which the bulk of TFIID dissociates from the promoter following the loading of TBP and early PIC formation. Thus, a function of TFIID might be to position and load TBP on the core promoter properly.

### BRG1 and SMARCB1 stimulate the formation of the fragile nucleosome -1 and its associated subnucleosomal particles at promoters

Acute inhibition of BRG1 activity causes a rapid and global loss of chromatin accessibility and transcription at promoters and enhancers in ES cells^32^. BAF complex activity is often assessed using the ATAC-seq method, which efficiently detects variations in chromatin accessibility^26,32^. However, the ATAC-seq signal does not discriminate the distinct components of the accessible chromatin that are fragile nucleosomes, the various subclasses of subnucleosomal particles, and histone-free TF-DNA complexes. To identify the functions of the BAF complexes in the control of the discrete nucleosomal and subnucleosomal organization described above, we examined how the loss of function of two essential BAF complex subunits impacts the histone H3 MNase ChIP-seq pattern at promoters and enhancers (Fig. 3). We selected for this analysis the BRG1 subunit, which is the main ATPase of all BAF complexes in ES cells^26^. We also chose SMARCB1, an essential subunit required for the interaction of cBAF and PBAF with their target nucleosomes; the cryoEM structure of the purified human cBAF complex revealed that SMARCB1 and BRG1 form a molecular clamp in which these subunits each interact with one of the two acidic patches on the nucleosome’s lateral faces, thus engaging the nucleosome bilaterally^33^. At promoters, 3h and 20h of BRG1 depletion reduced the production of 141-180 bp histone H3 ChIP fragments at position -1 to 75% and 50% of the signal detected in control cells, respectively (Fig. 3a, b and Extended Data Fig2a, b). This observation shows that BRG1 regulates the production of fragile nucleosomes at position -1. A 3h depletion of SMARCB1 resulted in a similar reduction (Fig. 3a-c). 3h depletions of BRG1 or SMARCB1 also decreased the production of 30-80 bp subnucleosomes to ∼65% of the signal detected in control cells (Fig. 3a, b and Extended Data Fig. 2d). In contrast, the production of 81-110 and 111-140 bp fragments was less affected by BRG1 or SMARCB1 depletion (Extended Data Fig. 2b, c).

**Figure 3.**
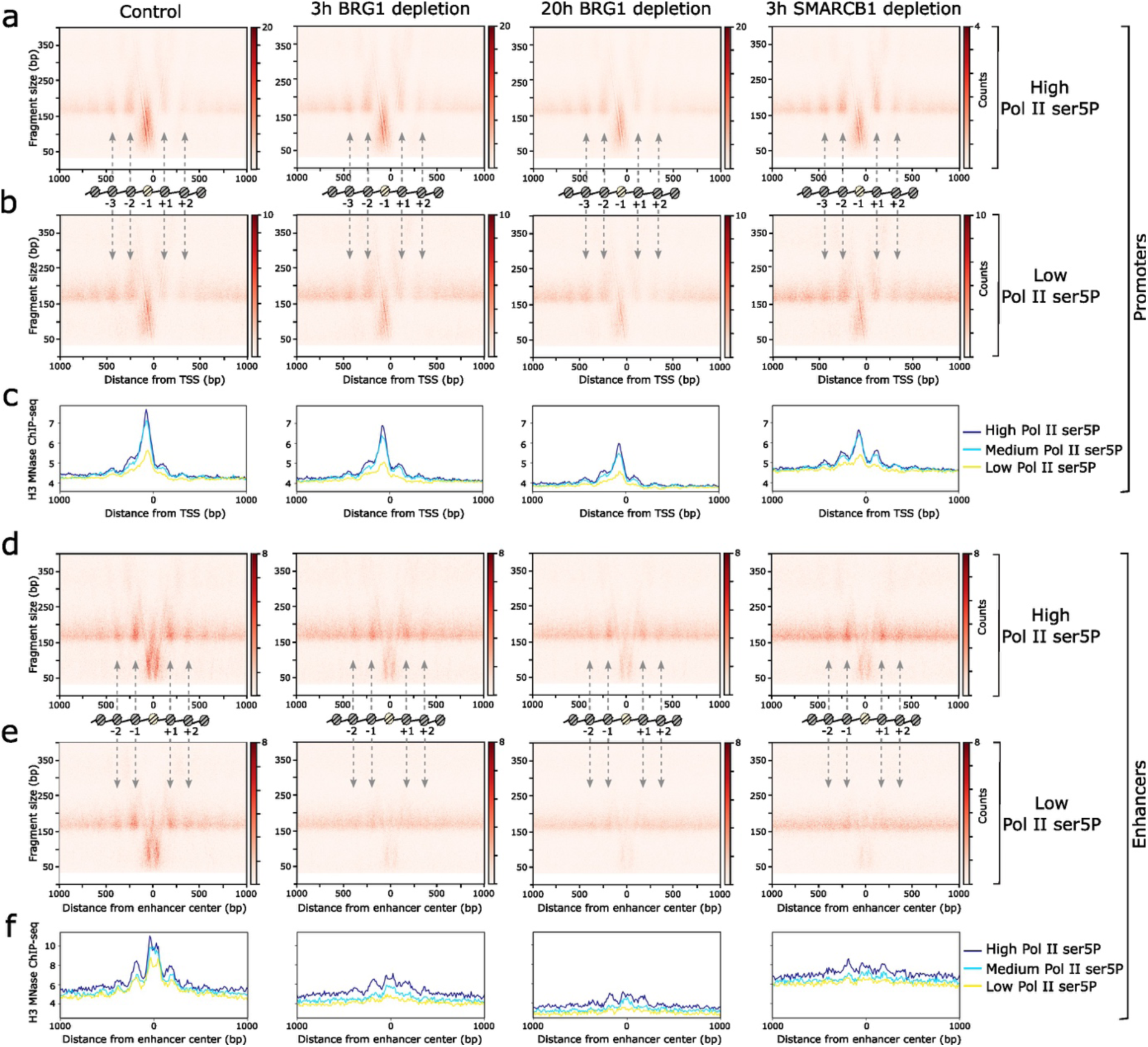
BRG1 and SMARCB1 differentially regulate the formation of the fragile nucleosome and subnucleosomal particles at promoters and enhancers. Chromatin was prepared from WT ES cells, or cells depleted of BRG1 or SMARCB1 for 3 or 20 h. Promoters and enhancers were divided in three subgroups according to Pol II-S5P ChIP-seq signal intensity (high, medium and low) **a, b**. V-plots of histone H3 ChIP-seq fragments spanning ±1,000 bp from the TSS of high (a) and low (b) Pol II-S5P promoters. **c**. Average H3 ChIP-seq profile of promoters subgroups sorted according to Pol II-S5P enrichment: high (dark blue), medium (light blue), or low (yellow). **d, e**. V-plots of histone H3 ChIP-seq fragments spanning ±1,000 bp from the enhancer center. The enhancers were sorted as above in subgroups according to Pol II-S5P occupancy. **f**. Average H3 ChIP-seq profiles of the enhancer subgroups. The schematic illustrations in a, b, d, e indicate the positions of the canonical (grey) and fragile (yellow) nucleosomes. At least two biological replicates were performed for each ChIP-seq experiment.

Pol II promoter-proximal pausing has been shown to stabilize BAF complex occupancy and may be a major determinant of BAF complex recruitment^22^. We thus separated in our analysis the promoters and enhancers in subgroups according to their level of enrichment for Pol II phosphorylated on serine 5 of the carboxy-terminal domain (hereby referred to as Pol II-S5P), which is the major isoform of paused Pol II. We observed that Pol II-S5P enrichment was correlated with the production of fragile nucleosomes at position -1: promoters of the low Pol II-S5P subclass produce less subnucleosomes than the medium and high subclasses (Fig. 3a, b and Extended Data Fig. 2a), suggesting that a Pol II-S5P-mediated recruitment of BRG1 might increase the generation of fragile nucleosomes. By contrast, the production of 30-80 bp subnucleosomes was not correlated with the level of Pol II-S5P (Extended Data Fig. 2d).

We compared the results of our histone H3 ChIP-seq analysis with the outcome of the analysis of BRG1 inhibition by the BRM014 molecule using the ATAC-seq approach^32^ (Extended Data Fig. 3a-c). The main ATAC-seq signal was identified in the genomic interval corresponding to the fragile nucleosome and its associated subnucleosomal particles (Extended Data Fig. 3a-b). A 2h BRM014-mediated inhibition of BRG1 decreased the ATAC-seq signal to ∼75% of its level in control cells (Extended Data Fig. 3c). This comparison of the alterations detected by histone H3 ChIP-seq and ATAC-seq show that, at promoters, the nucleosomal and subnucleosomal landscape and chromatin opening are affected in a subtle manner by BRG1 or SMARCB1 depletion.

### The control of nucleosomal and subnucleosomal organization by BRG1 and SMARCB1 reveals distinct involvements of BAF complexes at promoters and enhancers

At enhancers, 3h and 20h of BRG1 depletion reduced the production of fragile nucleosomes to only ∼20% of its level in control cells (Fig. 3d, e and Extended Data Fig. 2e). This acute reduction affected equally the Pol II-S5P-high, medium or low subclasses. In addition, depletion of BRG1 for 3 or 20h resulted in a severe alteration of nucleosomes -1 and +1 positioning (Fig. 3d, e and Extended Data Fig. 2e). This change in positioning was sensitive to the level of Pol II-S5P: nucleosomes -1 and +1 positioning was completely lost in the low Pol II-S5P enhancer subclass. While nucleosomal positioning was also severely impaired in the high Pol II-S5P subcategory, a pattern reminiscent of control cells subsisted (Extended Data Fig. 2e).

As previously described^21^, depletion of BRG1 for 3 or 20h also critically decreased the production of all subnucleosomal species at enhancers (Extended Data Fig. 2h). This decrease was correlated with the level of Pol II-S5P enrichment; the production of 30-80 bp subnucleosomes in the low Pol II-S5P enhancer subclass was reduced to ∼17% of the level detected in control cells, but the reduction was less extensive (34%) in the high Pol II-S5P subclass.

As above, we compared the results of our histone H3 MNase ChIP-seq-based analysis with the outcome of the ATAC-seq-mediated analysis of BRG1 function. In agreement with the situation at promoters, the main ATAC-seq signal was located in the genomic interval containing the fragile nucleosome and subnucleosomal particles (Extended Data Fig. 3d, e). BRM014-mediated inhibition of BRG1 resulted in a severe reduction of the ATAC-seq signal, mirroring the acute depletion of the subnucleosomal particles and fragile nucleosomes (Extended Data Fig. 3d-f). Our previous analysis using gradient MNase-seq revealed that the production of TF-DNA (histone-free)complexes was also strongly reduced in BRG1-depleted ES cells^21^.

In conclusion, BRG1 and SMARCB1 regulate the production of fragile nucleosomes and subnucleosomes at both promoters and enhancers, yet the situation is fundamentally different at these two categories of CRE. Enhancers are highly vulnerable to BRG1 inhibition or loss of function: each element of the discrete nucleosomal and subnucleosomal organization, and the TF-DNA complexes, are severely altered. Promoters are affected by a milder version of the same alterations; they conserve a proper positioning of canonical nucleosomes, and still produce (though at a reduced level) fragile nucleosomes, subnucleosomal particles and histone-free TF-DNA complexes.

### cBAF regulates early PIC formation at promoters and enhancers with different specificities

We uncovered above that the BAF complexes do not regulate promoters and enhancers to the same extent. Next, we wondered how the BAF complex would regulate early PIC formation at promoters and enhancers. For this purpose, we depleted key subunits of cBAF, PBAF, and ncBAF, and analyzed how the loss of function of each subunit would alter the binding of TBP and TFIIA to promoters and enhancers (Fig. 4). Among the subunits analyzed, BRG1 is the ATPase of all three complexes, SMARCB1 belongs to both cBAF and PBAF, whereas ARID1A, BRD7, and BRD9 are specific of cBAF, PBAF, and ncBAF, respectively. Intriguingly, ARID1A was the only subunit whose depletion resulted in a global reduction in the binding of TBP and TFIIA to promoters (Fig. 4a-d). Our analysis revealed that ARID1A stimulates the binding of TBP at ∼28% of the active promoters, and ∼20% of bivalent promoters. We explored whether the different subclasses of promoters based on the broad and sharp pattern of transcription initiation might be differentially regulated by ARID1A (Extended data Fig. 4). This analysis revealed that broad and sharp promoters are regulated to the same extent by ARID1A, irrespective of the presence of a TATA box (Extended data Fig. 4). Surprisingly, this stimulatory function of ARID1A is often independent of BRG1 and SMARCB1. Examination of individual genes revealed that only ∼8% of the active promoters and ∼12% of the bivalent category required BRG1 and SMARCB1 in addition to ARID1A to stimulate TBP binding to the core promoter. In sharp contrast, the binding of TBP and TFIIA to enhancers almost systematically required the activity of BRG1, SMARCB1, and ARID1A (Fig. 4e and Extended data Fig. 4).

**Figure 4.**
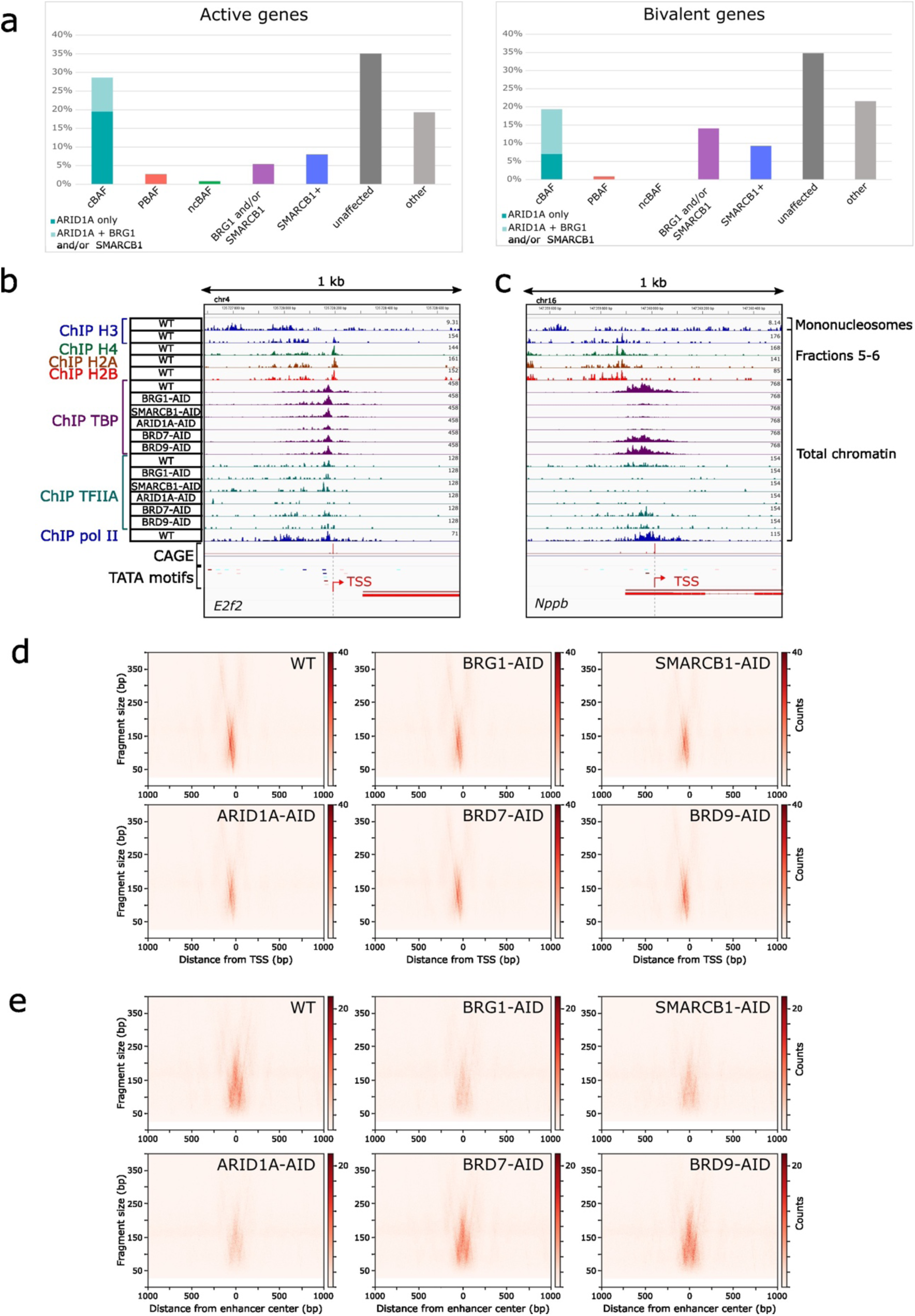
cBAF differentially regulates early PIC formation at promoters and enhancers. **a**. Histograms showing the percentage of active (left) or bivalent (right) promoters affected by the loss of function of BAF complex subunits, based on changes in TBP ChIP-seq signal. In the left panel, the promoters affected by the depletion of ARID1A alone are represented in dark turquoise, those affected by both ARID1A and BRG1, or both ARID1A and SMARCB1 depletion are in light turquoise. The genes affected by BRG1 and/or by SMARCB1 depletion, but not by ARID1A, are indicated in a different category. Additional bars represent the effects of the depletion of subunits of PBAF (BRD7 alone or in combination with BRG1 and/or SMARCB1, in pink) and ncBAF (BRD9 alone or with BRG1, in green). The “SMARCB1+” category (blue) corresponds to promoters where TBP binding is stimulated by SMARCB1 depletion. “Unaffected” (dark grey) includes promoters showing no change in TBP binding, and “other” (light grey) includes promoters affected by other combinations of depleted subunits. **b, c**. Density graphs showing the distribution of ChIP DNA fragment centers at representative examples of active (**b**) and bivalent (**c**) promoters. The top lane shows the positions of canonical nucleosomes detected by histone H3 ChIP-seq of sucrose gradient fractions 11 and 13. The next four lanes show the 50–80-bp subnucleosomes distribution, revealed by histone H3, H4, H2A and H2B ChIP-seq of sucrose gradient fractions 5 and 6. The subsequent lanes represent TBP, TFIIA, and Pol II ChIP-seq signal in WT cells or cells depleted for the indicated BAF subunits. CAGE signal maps the position(s) of the TSS. TBP consensus motifs with high (P = 0.001) and low (P = 0.01) confidence are indicated; the darker the color, the higher the confidence. For CAGE, TATA motifs and gene annotations, the blue and red colors indicate an orientation to the left and right, respectively. **d, e**. V-plots of TBP ChIP-seq fragments spanning from ±1,000 bp from the TSS for promoters (**d**) or the enhancer center (**e**). Chromatin was prepared from WT cells or cells depleted for the indicated subunits. At least two biological replicates were performed for each ChIP-seq experiment.

The extend to which the subunits of cBAF were required for TBP/TFIIA binding at promoters and enhancers was also drastically different. The binding of TBP was dependent on the three tested subunits of cBAF at a large majority of enhancers, whereas ARID1A alone or combined with BRG1 and SMARCB1 are required for TBP binding to < 30% of the active and <20% of the bivalent promoters (Fig. 4e and Extended data Fig. 4). This result is in frame with our analysis of cBAF involvement in regulating the nucleosomal and subnucleosomal landscape, which is extensive at enhancer and mild at promoters.

Our analysis also revealed that the depletion of subunits specific to PBAF and ncBAF complexes rarely affects TBP and TFIIA binding to the promoters or enhancers (Fig. 4a, d, e). PBAF and ncBAF are thus not involved in regulating early PIC formation at promoters or enhancers. Finally, our study also identified a subset (∼8%) of active and bivalent genes at the level of which the depletion of SMARCB1, but not of other BAF subunits, results in a stimulation of TBP binding at the promoter (Fig. 4a). RNA-seq analysis uncovered that 28% of these genes are upregulated by SMARCB1 depletion, suggesting that SMARCB1 has a repressive function targeting a subset of gene promoters.

### PBAF regulates nucleosomal and subnucleosomal organization at REST binding sites

Our data above show that BRG1 and SMARCB1 regulate chromatin organization and opening at both enhancers and promoters, though to different extents. Moreover, our analysis revealed that cBAF, but not PBAF or ncBAF, is required for early PIC formation at enhancers. At promoters, stimulation of TBP/TFIIA was often dependent on the ARID1A subunit, but not on subunits specific of PBAF or ncBAF, again pointing to a regulatory function exclusively carried out by cBAF. These results suggest that in ES cells, PBAF and ncBAF complexes are dispensable for chromatin remodeling at enhancers and promoters. To investigate in an unbiased manner whether BRD7, PBRM1, and BRD9 might be required to regulate the binding of TF onto the mouse genome, we analyzed the TF DNA motifs present in the DNA prepared from the top sucrose gradient fractions after centrifugation of MNase-digested chromatin from BRD7-, PBRM1-, and -BRD9-depleted ES cells. The top sucrose gradient fractions contain the TF-DNA complexes released by MNase digestion from CRE (REF^21^ and Fig. 2). In parallel, we also similarly analyzed the TF motifs detected from BRG1-, SMARCB1-, ARID1A-depleted, and control cells. This analysis identified a severe depletion of TF motifs such as OCT4/SOX2/NANOG, which are associated to enhancer function, in BRG1-, SMARCB1-, and ARID1A-depleted cells (Fig. 5a).

**Figure 5.**
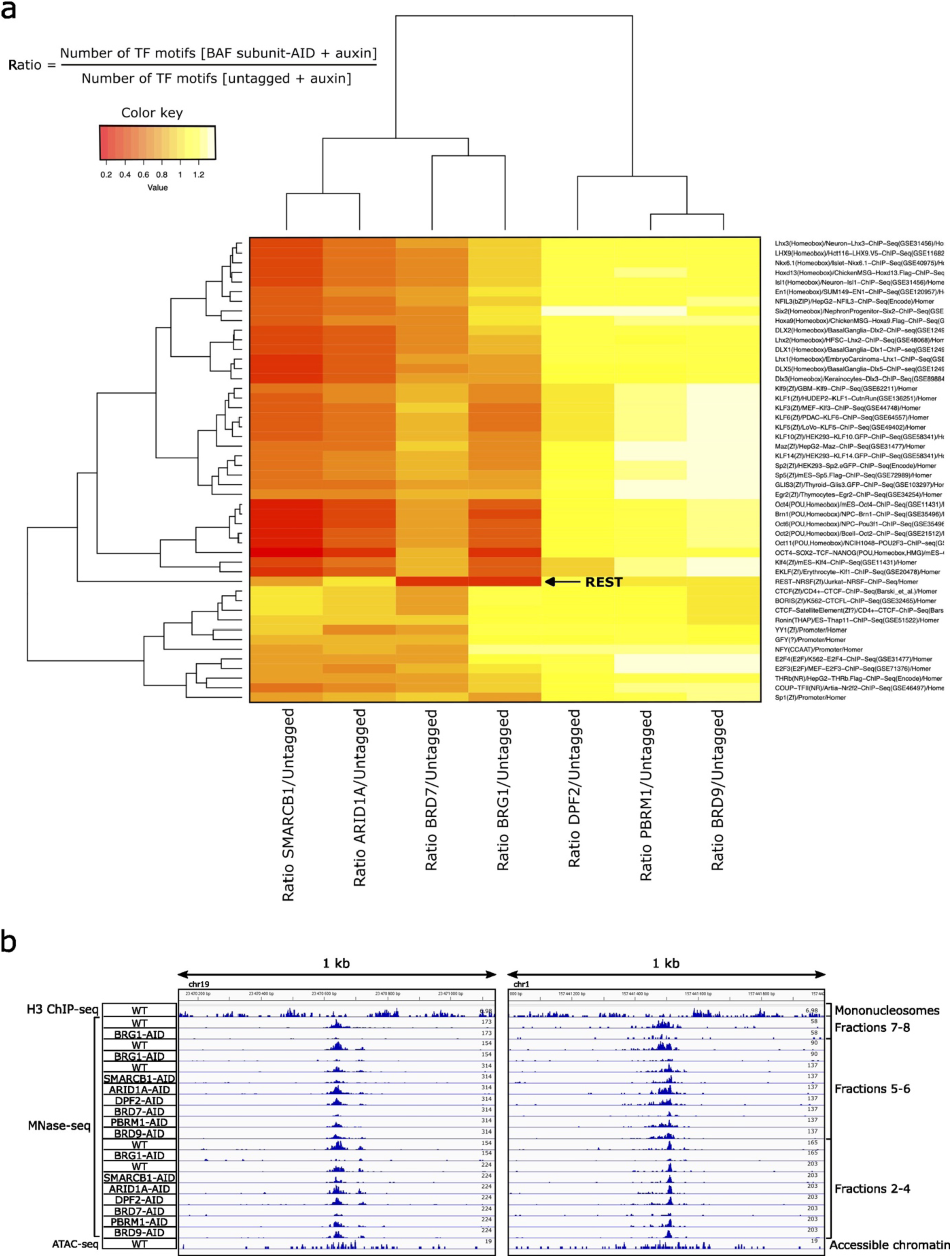
PBAF regulates nucleosomal and subnucleosomal organization at REST binding sites. MNase-digested chromatin, prepared from control ES cells, or from cells depleted of BRG1, SMARCB1, ARID1A, DPF2, BRD7, PBRM1, and BRD9 using the AID system, was centrifugated through a sucrose gradient as in Fig. 2. **a**, The TF motifs present in the DNA prepared from the pool of sucrose gradient fractions 2-4, containing the CRE-associated TF-DNA complexes, were detected using the HOMER tool. The ratio between the number of TF motifs detected in the BAF subunit-depleted versus control samples was calculated and visualized as a heatmap. **b**, Density graphs showing the distribution of DNA fragment centers at representative examples of REST binding sites. The top lane shows the positions of canonical nucleosomes detected by histone H3 ChIP-seq of sucrose gradient fractions 11 and 13. All the following lanes except the last represent gradient MNase-seq data (the DNA from the pools of sucrose gradient fractions was sequenced without any ChIP). The first two lanes following the mononucleosomes represent sucrose gradient fractions 7-8 in the control or BRG1-depleted conditions. The following nine lanes represent fractions 5-6 in control cells or cells depleted for the indicated BAF subunits for 20 h. The following nine lanes represent fractions 2-4 in the same contexts. The last lane represents accessible chromatin mapped by ATAC-seq in WT ES cells.

Interestingly, BRD7-, PBRM1- or BRD9- depletion had little effects on these enhancers-associated motifs, suggesting that PBAF and ncBAF do not regulate the binding of TFs to enhancers. Our previous analysis had revealed that BRG1 does not regulate CTCF binding to its DNA motifs^21^. In agreement with this result, we did not detect any significative alteration of CTCF binding in ARID1A-, BRD7-, PBRM1- or BRD9-depleted cells (Fig. 5a). We concluded that none of the BAF complexes is involved in chromatin remodeling events affecting CTCF binding to DNA.

In contrast to the situation observed at CTCF site, our analysis revealed that the depletion of both BRD7 and BRG1 resulted in a marked depletion of the REST binding motifs, providing evidence that the PBAF complex regulates chromatin opening at REST sites (Fig. 5a). Examination of individual REST binding sites confirmed the critical requirement for BRD7 and BRG1 for the production of histone-free REST- DNA complexes. In addition, BRD7 and BRG1 were also required for the production of subnucleosomal particles and fragile nucleosomes at REST sites (Fig. 5b and Extended Data Fig. 5). Surprisingly, at most REST binding sites, the PBRM1 subunit of PBAF was dispensable for both the generation of histone-free REST-DNA complexes and subnucleosomes, while SMARCB1 had only a minor role. This result demonstrates that not all subunits of the PBAF complex are required to remodel the chromatin of REST binding sites.

## DISCUSSION

In this study, we have defined the nucleosomal and subnucleosomal organization of the core promoter region in mouse embryonic stem (ES) cells, and examined how TBP and TFIIA, which are the first GTFs initiating PIC formation, interact with this chromatin landscape. We have separated the core promoters in distinct subgroups (broad and sharp) according to the shape of transcription initiation and the presence or absence of a TATA box. We observed that early PIC formation occurs in the same region of the core promoter in which the fragile nucleosomes and subnucleosomal particles are located. Our data show that PIC formation can follow two distinct pathways. At TATA-less promoters, which correspond to the most abundant subclasses of broad and sharp promoters, PIC formation occurs onto a genomic template mostly depleted of histones. In contrast, at TATA box-containing promoters, PIC formation is initiated in the context of a core promoter still partially wrapped into a fragile nucleosome or a subnucleosomal particle. This suggests that *in vivo*, the binding of GTFs to TATA-box containing promoters involves interactions with both DNA and histones. This second pathway might provide a supplementary constraint on the binding of the GTFs to the core promoter, helping to focus transcription initiation.

We observed a similar situation at enhancers, which in their large majority do not contain a TATA-box. At TATA-less enhancers, the interaction of TBP and TFIIA occurs predominantly in the context of a template devoid of histones. As for promoters, the presence of a TATA box within the enhancer was generally associated with the co-enrichment of TBP with histones H3 and H2B.

The formation of the PIC begins with TFIID binding to the core promoter via the TATA box and/or DPE, triggering structural changes in TFIID that position TBP on the core promoter. After these rearrangements, TBP remains connected to TFIID solely through TFIIA^10–12^. It is unclear if this new conformation remains stable during the subsequent steps of PIC formation, or if it is the prelude to dissociating the rest of TFIID from the core promoter. Our combination of sucrose gradient centrifugation and ChIP-seq experiments show that TBP is present at promoters in complexes having a molecular size much smaller than TFIID, suggesting that TBP can remain at the promoter without the complete TFIID complex. This supports a model in which one of TFIID’s primary roles is to position and load TBP during early PIC formation. An alternative scenario is that TBP could bind the TATA-box without being a subunit of TFIID.

We have analyzed how the BAF complexes regulate early PIC formation at core promoters and enhancers. We observed that the cBAF complex, but not PBAF and ncBAF, regulates PIC formation at these CRE. However, the importance of cBAF in this process is different at enhancers and promoters. The ARID1A subunit of cBAF stimulates the binding of TBP and TFIIA at a subset of promoters. In contrast, all tested subunits of cBAF are required to allow the binding of TBP and TFIIA at enhancers. This differential control of PIC formation agrees with a greater involvement of cBAF in organizing the nucleosomal and subnuclesomal organization of enhancers.

Our investigations also uncovered that PBAF and ncBAF do not control the chromatin organization or opening at promoters and enhancers in ES cells. Previous studies based on the binding profile of subunits of PBAF and ncBAF suggested that PBAF and ncBAF might be required for chromatin remodeling at promoter and CTCF binding sites^34–36^. Our study in ES cells do not confirm these hypotheses. Based on the analysis of TF motifs present in the CRE-associated DNA-TF complexes purified by sucrose gradient centrifugation, we detected a highly specific function of PBAF in regulating the nucleosomal and subnucleosomal landscape of REST binding sites. Interestingly, not all subunits of PBAF were equally involved in this process. The BRD7 and BRG1 subunits were required to generate subnucleosomes and TF-DNA complexes at more than 75% of the REST sites, but PBRM1 was dispensable. The SMARCB1 subunit, based on the structural model of cBAF and PBAF, is required with BRG1 to interact with its target nucleosomes. We would thus expect SMARCB1 to be as essential as BRG1 for proper chromatin remodeling at REST sites. However, SMARCB1 was required to produce subnucleosomes and TF-DNA complexes only at ∼20% of the REST sites. These differences in requirement between BRG1, BRD7, SMARCB1, and PBRM1 were not due to a lesser efficiency of the auxin-degron system for degrading each subunit. This analysis suggests that the different subunits of the PBAF complex each have specific functions that are not necessarily required at all PBAF REST target loci. Our results agree with a recent publication showing that PBAF facilitates the binding of REST to its target loci^37^.

Our analysis of cBAF function at promoters and enhancers also points to specific functions for the different subunits of this complex. As for PBAF, the requirement for each subunit may depend on the context of each promoter or enhancer. At promoters, we observed that ARID1A is required to stimulate the binding of TBP, and this activity was often independent of BRG1 and SMARCB1 activities. This ARID1A-dominated regulation at promoters suggests that cBAF does not require its nucleosome binding function, nor its DNA translocase activity to stimulate the binding of TBP. ARID1A has been identified as a key scaffold protein involved in the assembly of the cBAF complex^31^. This work revealed a new function of ARID1A in recruiting the GTF TBP to the core promoter.

Our study also revealed that BRG1 or SMARCB1 loss-of-function preferentially alters chromatin remodeling at enhancers negative for Pol II-S5P and total Pol II. Following the degradation of BRG1 or SMARCB1, the nucleosomal and subnucleosomal organization of Pol II-S5P-high enhancers is less extensively altered than at Pol II-S5P-low enhancers. In contrast, the BRM014 molecule, which inhibits BRG1, similarly alters the ATAC-seq signal at both categories, suggesting that Pol II-S5P-high and -low enhancers are equally dependent on BRG1. A possible explanation for this difference is that the inhibitor targets all BRG1 molecules, whereas the auxin-mediated degradation of BRG1- and SMARCB1-AID results in a low level of residual protein^21^. The BAF complexes that can still assemble from these residual proteins might be preferentially targeted to Pol II-S5P-rich enhancers according to the mechanism described in reference^22^. This mechanism preferentially targeting residual BAF complexes to active, Pol II-S5P-enriched enhancers has important consequences for enhancer function during cell differentiation. In ES cells and many precursor cell types, enhancers can harbor three main different statuses: active, primed (or poised), and silent^38,39^. Active enhancers regulate gene expression in ES cells, whereas primed enhancers are not active, but are committed to becoming active during differentiation. Once differentiation initiates, primed enhancers will switch to the active status and begin to regulate gene expression. In ES cells, the active and primed enhancers are both bound by the pluripotency-associated TFs, including OCT4, SOX2 and NANOG, that maintain the ES cell phenotype^40^. What distinguishes the primed from the active enhancers is their transcription status, and primed enhancers can be identified based on their low Pol II-S5P enrichment. Our data show that primed enhancers will be the most altered upon BRG1 or SMARCB1 loss of function. This extensive alteration of primed enhancer organization will jeopardize enhancer function during differentiation, and therefore alter the gene expression pathways controlling cell differentiation. This mechanism is likely to contribute to the transformation events triggered by SMARCB1 loss of function mutations during development, leading to rhabdoid tumors arising shortly after birth or during early childhood^30,41^.

In conclusion, our study allowed us to define better the chromatin remodeling functions of the cBAF and PBAF complexes. The activity of mammalian BAF complexes had been previously extensively studied by focusing on their chromatin “opening” function, defined by the ATAC-seq approach. The ATAC-seq signal has generally been interpreted as histone-free DNA generated by the remodelers to allow the binding of the TF targeting their motif on each CRE. Little information was available on how the BAF complexes regulate the production of fragile nucleosomes and subnucleosomal particles. In this comprehensive study, we have combined the results of several experimental approaches to revisit and clarify the function of the BAF complex at CRE. Our integrative study identified specific functions of cBAF and PBAF’s subunits in controlling TF binding to CRE, and their nucleosomal and subnucleosomal organization.

## SUPPLEMENTARY FIGURES

**Extended Data Fig. 1.**
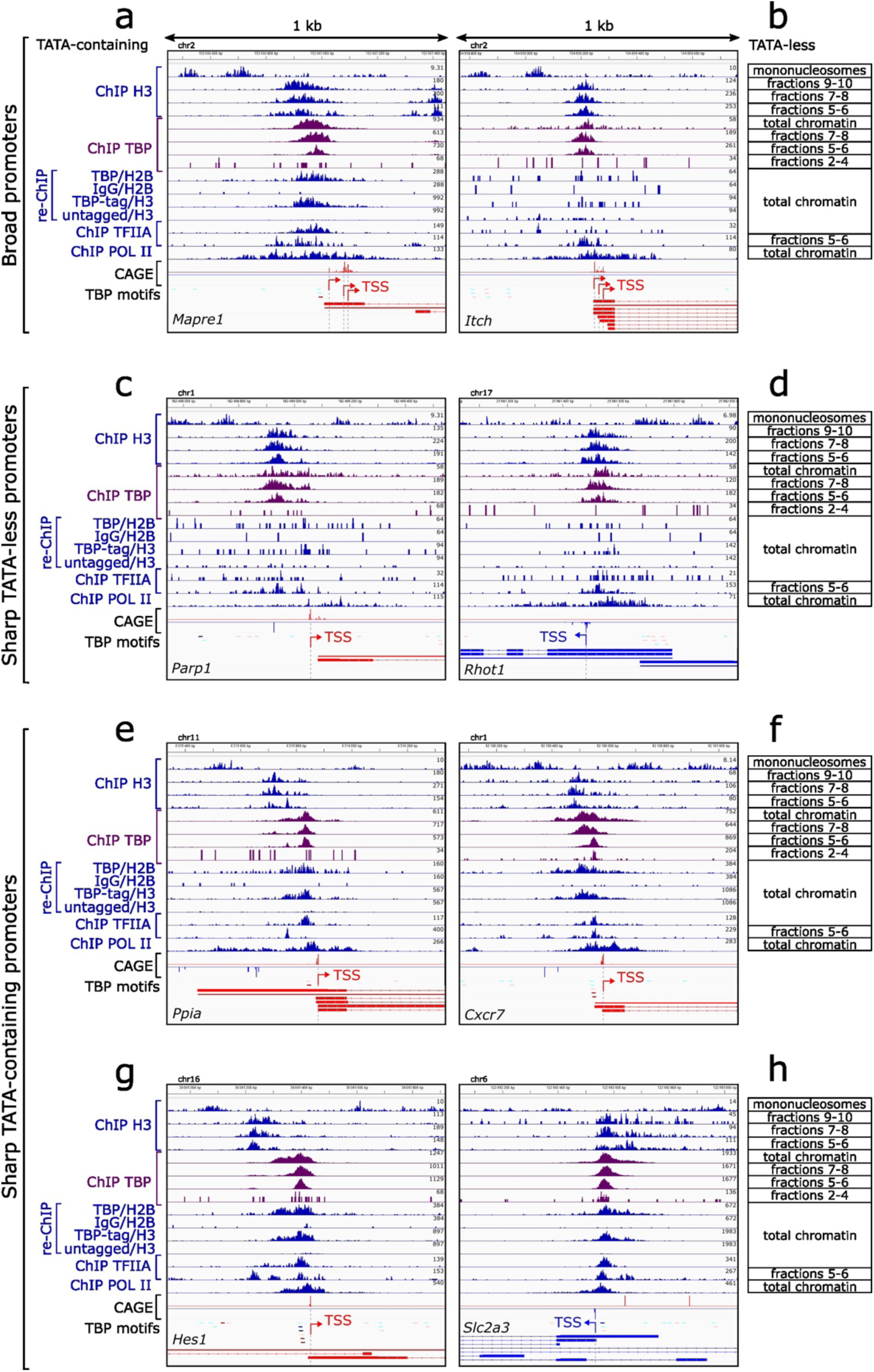
TBP binds the TATA box in the context of a histone-containing subnucleosomal particle. Density graphs showing the distribution of ChIP DNA fragment centers at representative examples of the three types of promoters, as indicated on the left. The top lane shows the positions of canonical nucleosomes detected by histone H3 ChIP-seq of sucrose gradient fractions 11 and 13. The three following lanes show the position of the fragile nucleosomes (fractions 9-10) and the different subnucleosomal species (fractions 7-8 and 5-6), revealed by histone H3 ChIP-seq of sucrose gradient fractions (see Figure 2). The following lanes show the TBP ChIP-seq signal using as input either total chromatin (MNase-digested chromatin, unseparated on a sucrose gradient), or subnucleosomal fractions 7-8 and 5-6, and histone-free TF-DNA fractions 2-4. The four subsequent lanes represent sequential ChIP-seq signal corresponding respectively to Figure 1 (c), (d), (e) and (f). Two lanes represent TFIIA ChIP-seq signal on total chromatin and fractions 5-6. The subsequent lane represents Pol II ChIP-seq signal using total chromatin as input. CAGE signal maps the position of the TSS. TBP consensus motifs with high (P = 0.001) and low (P = 0.01) confidence are indicated, the darker the color, the higher the confidence. The last lane corresponds to gene annotations. For CAGE, TATA motifs and annotations, the color corresponds to the gene orientation: the genes depicted in blue are oriented to the left, while the genes depicted in red are oriented to the right. The TSS is symbolized by a unique arrow for sharp promoters and multiple arrows for broad promoters. At least two biological replicates were performed for each ChIP-seq experiment.

**Extended Data Figure 2.**
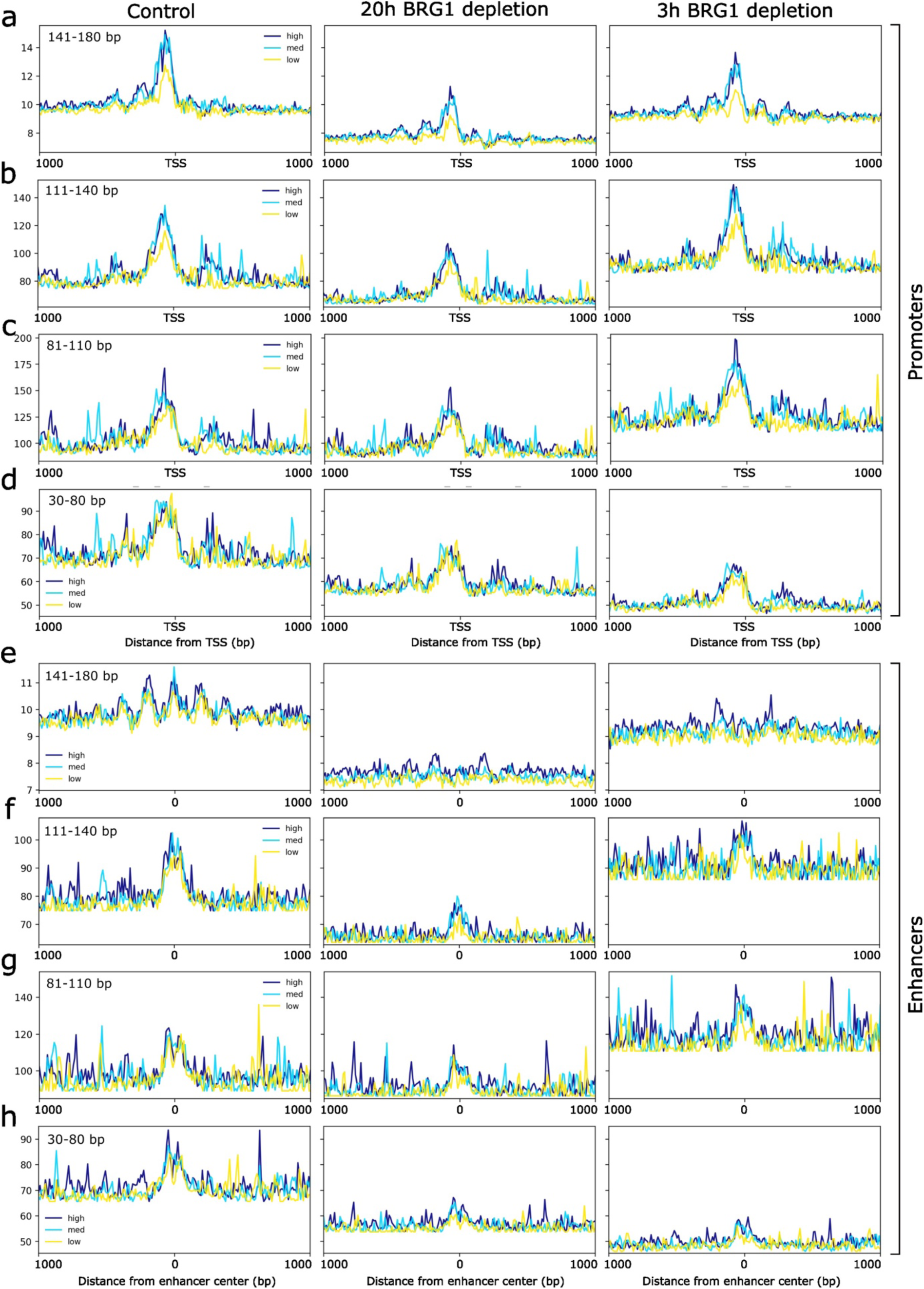
Nucleosomal and subnucleosomal species exhibit differential sensitivity to BRG1-mediated remodeling at promoters and enhancers. Average H3 ChIP-seq profiles at promoters (**a-d**) and enhancers (**e-h**), centerd on the TSS or the enhancer center, regarding the Pol II-S5P occupancy: high Pol II-S5P (dark blue), medium Pol II-S5P (light blue), or low Pol II-S5P (yellow). For all panels, chromatin was prepared either from wild-type (WT) cells, or from BRG1-depleted cells for 20 or 3 h; the condition is indicated on the top. The fragments are separated according to their size: 141-180 bp (**a, e**), 111-140 bp (**b, f**), 81-110 bp (**c, g**) or 30-80 bp (**d, h**).

**Extended Data Figure 3.**
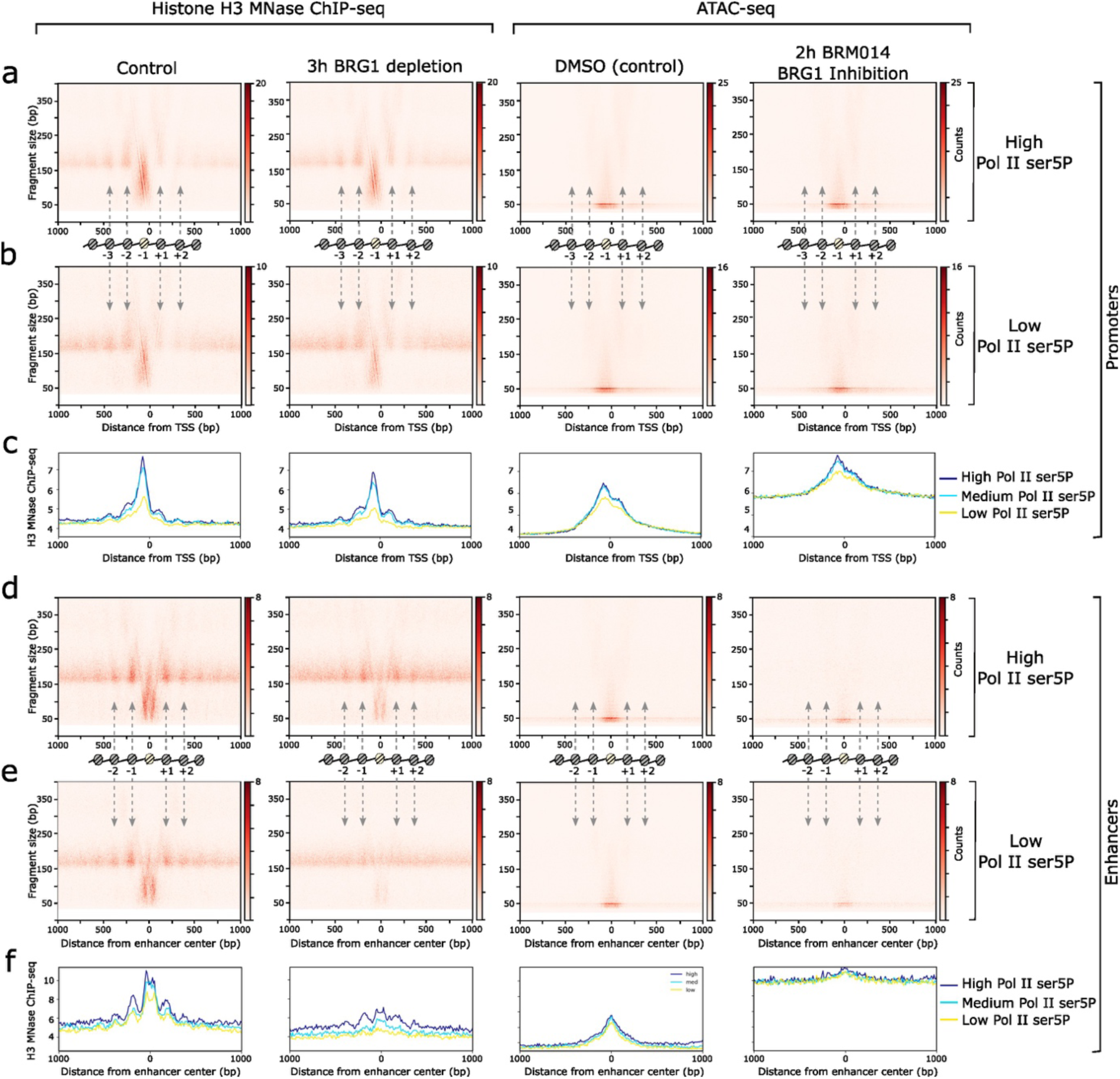
Comparison of the MNase ChIP-seq and ATAC-seq approaches to analyze chromatin remodeling by BRG1 at promoters and enhancers. For the whole figure, the two columns on the left side show the H3 ChIP-seq signal, while the two columns on the right side represent the ATAC-seq enrichment. For H3 ChIP-seq, the chromatin was prepared from WT (first column) or BRG1-depleted (second column) cells. For ATAC-seq, the chromatin was prepared from WT cells in the control condition (third column) or BRM014-treated condition (fourth column). Promoters are represented in panels **a-c**, while enhancers are represented in panels (**d-f**). **a, b**. V-plots of H3 ChIP-seq and ATAC-seq fragments spanning ±1,000 bp from the TSS. The promoters are sorted regarding the Pol II-S5P occupancy: high (a) vs low (b) Pol II-S5P. **c**. Average H3 ChIP-seq and ATAC-seq profiles centered on the TSS, regarding the Pol II-S5P occupancy: high (dark blue), medium (light blue), or low (yellow) Pol II-S5P. **d, e**. V-plots of histone H3 ChIP-seq and ATAC-seq fragments spanning ±1,000 bp from the enhancer center. The enhancers are sorted regarding the Pol II-S5P occupancy: high (d) vs low (e) Pol II-S5P. **f.** Average H3 ChIP-seq and ATAC-seq profiles centerd on the enhancer center, regarding the Pol II-S5P occupancy: high (dark blue), medium (light blue), or low (yellow) Pol II-S5P. The schematic illustrations in a, b, d, e indicate the positions of the canonical (grey) and fragile (yellow) nucleosomes.

**Extended Data Figure 4.**
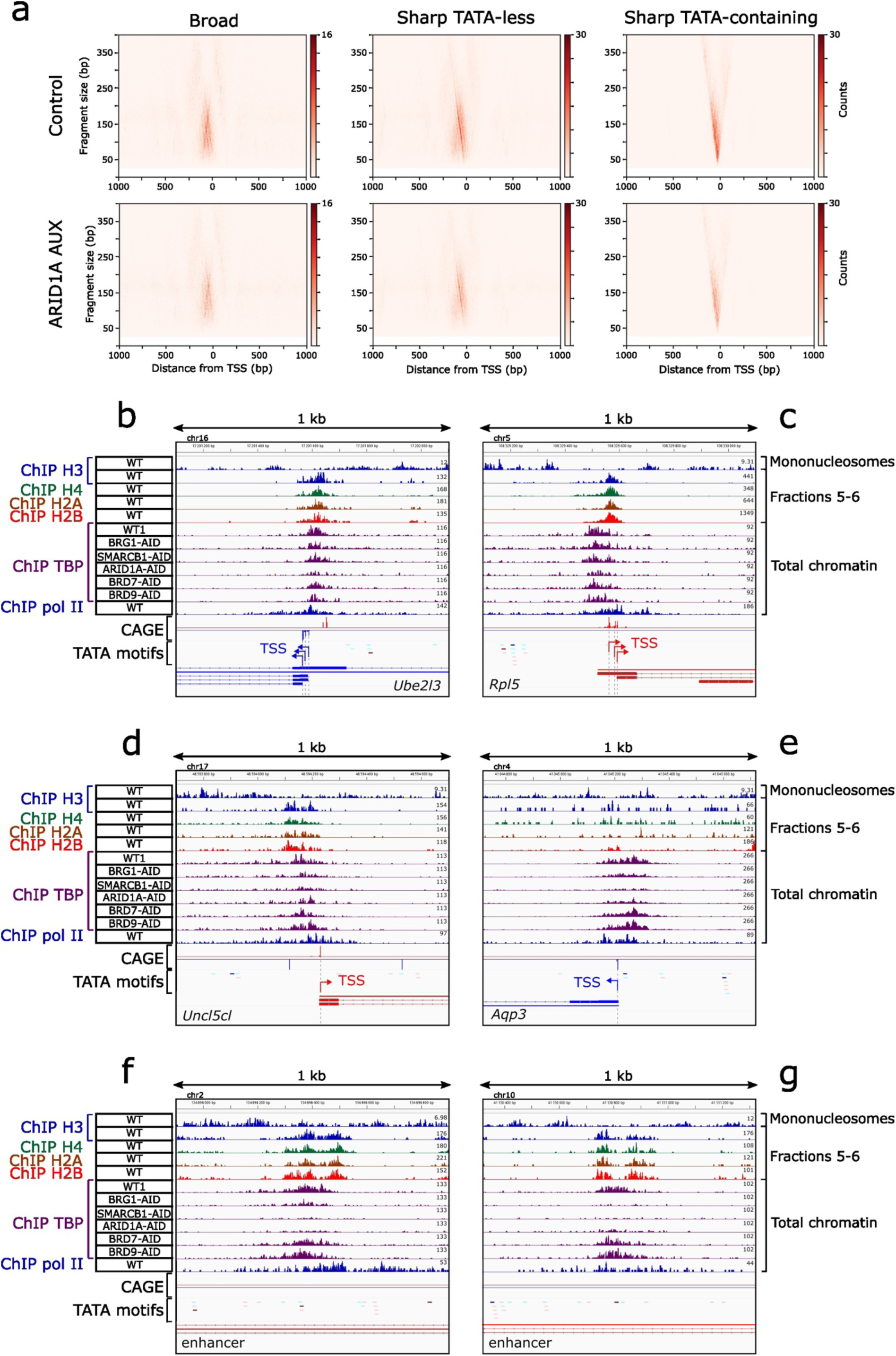
associated with Figure 4 – cBAF regulates PIC formation at all subclasses of promoters. **a**. V-plots of TBP ChIP-seq fragments spanning from ±1,000 bp from the TSS for promoters, in the WT cells (top) or ARID1A-depleted cells (bottom). The promoters are sorted according to their type, which is indicated at the top. **b-g**. Density graphs showing the distribution of ChIP DNA fragment centers at representative examples of active (**b, c**) and bivalent (**d, e**) promoters, and enhancers (**f, g**). The top lane shows the positions of canonical nucleosomes detected by histone H3 ChIP-seq of sucrose gradient fractions 11 and 13. The next four lanes show the 50–80-bp subnucleosomes distribution, revealed by histone H3, H4, H2A and H2B ChIP-seq of sucrose gradient fractions 5 and 6. The six subsequent lanes represent TBP ChIP-seq signal in WT cells or cells depleted the indicated BAF subunits. The following lane represents Pol II ChIP-seq signal. CAGE data are represented to map the position of the TSS. TBP consensus motifs with high (P = 0.001) and low (P = 0.01) confidence are indicated; the darker the color, the higher the confidence. The last lane corresponds to gene annotations. For CAGE, TATA motifs and annotations, the color corresponds to the gene orientation: the genes depicted in blue are oriented to the left, while the genes depicted in red are oriented to the right. The TSS is indicated by a unique arrow for sharp promoters and multiple arrows for broad promoters. At least two biological replicates were performed for each ChIP-seq experiment.

**Extended Data Figure 5.**
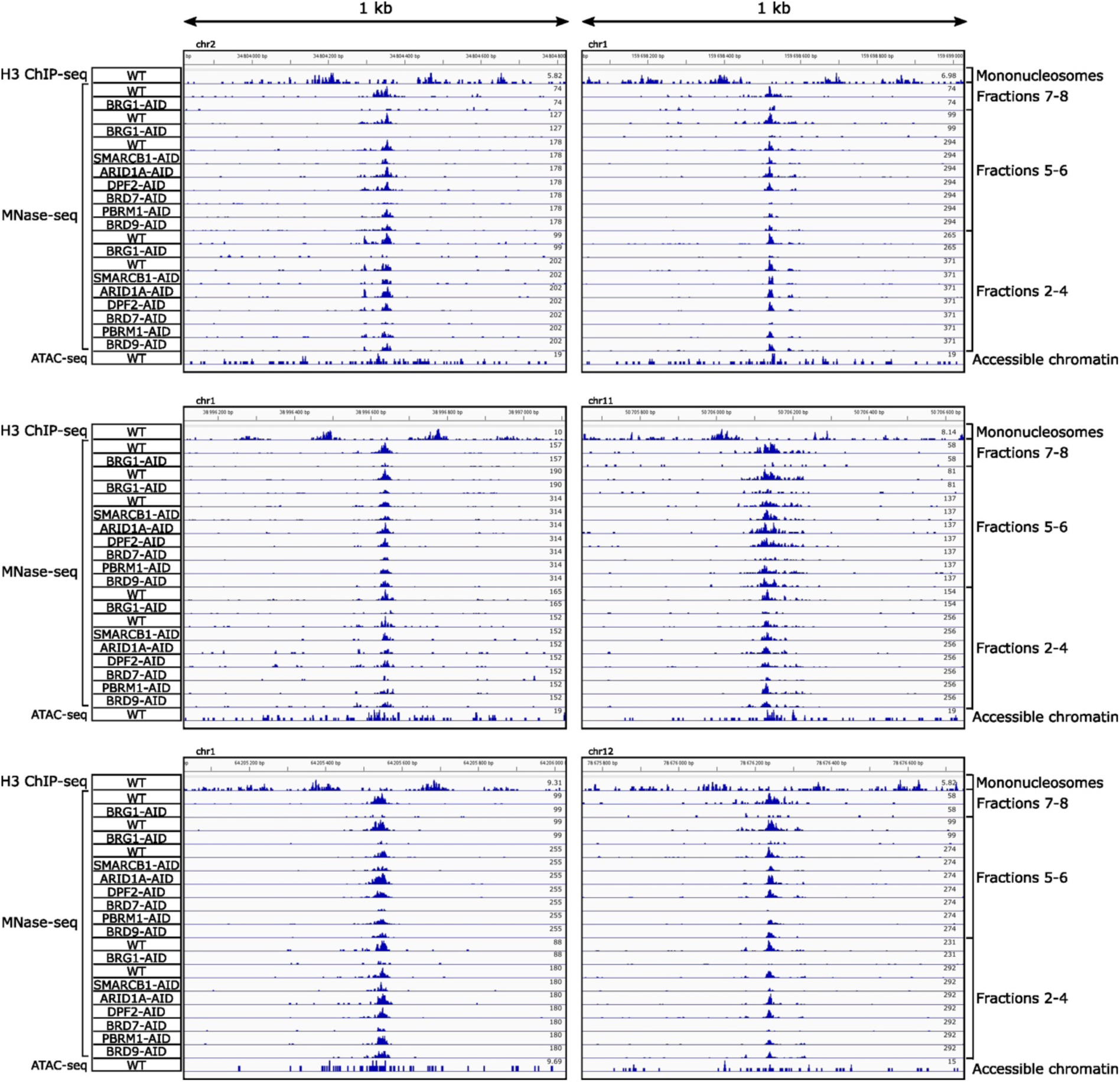
associated with Figure 5 – Additional examples of how PBAF regulates nucleosomal and subnucleosomal organization at REST binding sites. Density graphs showing the distribution of DNA fragment centers at representative examples of REST binding sites. The top lane shows the positions of canonical nucleosomes detected by histone H3 ChIP-seq of sucrose gradient fractions 11 and 13. All the following lanes until the penultimate represent gradient MNase-seq data (gradient fractions were sequenced without any ChIP). The first two lanes following the mononucleosomes represent sucrose gradient fractions 7-8 in the control or BRG1-depleted conditions. The following nine lanes represent fractions 5-6 in control cells or cells depleted for the indicated BAF subunits for 20 h. The following nine lanes represent fractions 2-4 in the same context as fractions 5-6. The last lane represents accessible chromatin mapped by ATAC-seq. At least two biological replicates were performed for each ChIP-seq experiment.

## METHODS

### Cell lines and culture

All mouse ES cell lines were grown on gelatinized cell culture dishes at 37 °C, 5 % CO2, in DMEM (Sigma) supplemented with leukemia inhibitory factor, 1X non-essential amino acids (Invitrogen), 15 % fetal calf serum (Invitrogen), 0.2 % β-mercaptoethanol (Sigma) and 1X penicillin/streptomycin (Invitrogen). The mAID-tagged cell lines were generated as previously described^21^. The E14Tg2a-Tir1 cell line was obtained from E. Nora and B. Bruneau^42^. The TBP-3FTH-mAID was generated following the same strategy as the cell lines previously described^21^. The sequences of the oligonucleotides encoding the sgRNAs targeting *Tbp* are :

Tbp_SG1_up : **CACCG**GAACATGGCAGACAACTATG; Tbp_SG1_down : **AAAC**CATAGTTGTCTGCCATGTTC**C**; Tbp_SG2_up : **CACCG**ATAGGGAAGGCAGGAGAACA; Tbp_SG2_down : **AAAC**TGTTCTCCTGCCTTCCCTAT**C**

### Auxin-mediated depletion of BAF subunits

Auxin-mediated protein depletion was performed as previously described^21^. Depletion was conducted during 20h or 3h. The depletion of the tagged protein was confirmed by western blotting.

### Chromatin preparation from formaldehyde-fixed cells

Chromatin preparation from formaldehyde-fixed cells was performed as described in reference^21^, with the exception of the MNase dose, which was set at 3 Kunitz units of MNase (New England Biolabs, 200 Kunitz units per µl) per 1 million cells.

**Separation of nucleosomes and subnucleosomal particles by sucrose gradient centrigugation** Fractionation of MNase-digested chromatin by sucrose gradient centrifugation was performed as described in reference^21^.

### ChIP-seq and sequential ChIP-seq (reChIP)

MNase ChIP-seq using formaldehyde-fixed chromatin was performed as described in reference^21^. We used two distinct sequential ChIP-seq protocols to test the co-association of TBP and histones. In the first protocol, we used antibodies against endogenous TBP and histone H2B. We prepared the chromatin from 120 million E14Tg2a mouse ES cells for each experiment. We incubated the chromatin with either 30 µl of monoclonal antibodies against TBP (G01092, Bertin Bioreagent) or 60 µg of nonspecific immunoglobulin G (IgG, I5381, Sigma). After a 16 h at 4 °C onto a rotative agitator, we added protein G agarose beads to chromatin–antibody complexes and further incubated for 4 h at 4°C. After three washes with TEN buffer (20 mM Tris-HCl pH 7.5, 150 mM NaCl, 3 mM MgCl2, 0.1 mM EDTA, 0.01% Igepal) and four washes with WBLiCl buffer (50 mM Hepes pH 7.5, 500 mM LiCl, 1 mM EDTA, 1 % IGEPAL, 0.7 % Na-deoxycholate), we incubated the beads with TEN buffer without Igepal containing 1 µg/µL TBP elution peptide (MDQNNSLPPYAQGLASP). This peptide, which corresponds to the first 17 aa of TBP, had been used for the production of the monoclonal antibody^43^. We performed three consecutive elution steps: 2 h at room temperature, 16 h at 4°C and 2 h at room temperature. The three elution fractions were pooled and incubated with 5 µg of antibodies against histone H2B (Abcam, ab1790) for 16 h at 4 °C. Protein G agarose beads were next added and incubated for 4 h at 4°C. We performed the wash and elution steps of this second ChIP as described above for regular ChIP-seq experiments. Two biological replicates were obtained for each TBP and nonspecific IgG antibody. In the second protocol, we used antibodies against the HA tag of TBP-3FTH and histone H3. For each experiment, we prepared the chromatin from 120 million TBP-3FTH-mAID-tagged mouse ES cells. We incubated this chromatin with 150 µL of agarose beads coupled to a mouse monoclonal anti-HA antibody (A2095, Sigma). As a control, the same experiment was realized using chromatin prepared from E14Tg2a mouse ES cells. After overnight incubation at 4°C, the beads were washed three times with TEN buffer and four times with WBLiCl buffer. The beads were resuspended in TEEP80 buffer and the elution was performed by adding 60 units of TEV protease (P8112S, New England Biolabs) during 90 min at 30°C. The eluate was next incubated with 10 µg of antibodies against histone H3 (Abcam ab1791) for 16 h at 4 °C. Protein G agarose beads were next added and incubated for 4 h at 4 °C. We performed the wash and elution steps of this second ChIP as described above for regular ChIP-seq experiments. Two biological replicates were obtained for each condition.

Table of antibodies used for ChIP-seq.

| Antibody | Reference | Clone | Quantity/ChIP |
| --- | --- | --- | --- |
| **H2B** | Abcam ab1790 | Polyclonal | 5 µg |
| **H3** | Abcam ab1791 | Polyclonal | 5 µg |
| **Pol II** | Santa Cruz sc55492 | F12 | 20 µg |
| **TFIIA-γ** | Santa Cruz sc374483 | D6 | 10 µg |
| **TBP** | Antibodies online mab60497 | 1TBP18 | 10 µg |
|  | Invitrogen MA1-10883 | 1TBP18 | 10 µg |
|  | Bertin Bioreagent G01092 | 3G3 | 10 µL |

### Computational analyses

#### Lists of sharp and broad promoters

Promoters transcribed in mouse ES cells were initially rank-ordered according to the distance between the two divergent TSSs located on the plus and minus strands, using 5’ ends of Start-RNA reads (GSE43390), as described^21^. The promoters were then separated in 6 groups of ∼1,000 genes, named G0-G5 from the shortest to largest distance. We then classified the genes in each of these groups according to the shape of their TSS into “sharp” and “broad” subgroups. We used CAGE-seq signal (GSM3188172) to determine the sharp and broad nature of each promoter. The coverage of the first six bases of the 5’ read extremities were separated on the minus and the plus strand. For each strand, we performed a peak calling using Macs2. Peaks with less than five reads were discarded. When a peak was unique in a window of 1,000 paired bases around the TSS annotation, and if the peak was under 5 bases in width, and if the signal had the same orientation as the annotation, the TSS was classified in the sharp group. When the peak’s width was larger than 5 bases, or when we detected multiples peaks within the 1000 bases window, the TSS was classified in the broad TSS group. Each TSS was manually verified and, when necessary, its coordinates were adjusted on the genome based on the CAGE-seq coverage profiles observed on the IGV browser. For sharp TSS, the base chosen for the new start site coordinate was the base with the highest CAGE-seq signal within the peak.

### Lists of TATA-containing promoters

Using Fimo, from the MEME suite^44^, we located the potential TBP motifs within a window of +/- 1000 bases around the TSS. Motifs with a P value lower than 0.001 were labeled as high confidence TBP motif. The TSS was considered as “TATA-box containing” when the distance between the TSS and the TBP motif was lower than 50 bases.

### Sorting promoters and enhancers according to Pol II-S5P ChIP-seq signal

The enhancer list used for Pol II-S5P ranking is the fusion of ES cell enhancer clusters 1 and 8 from reference^21^. The promoter list used for Pol II-S5P ranking is the fusion of the G1-G2 lists described above. We used computeMatrix to sort these promoter or enhancer list according to the decreasing Pol II-S5P coverage. Each Pol II-S5P-rank-ordered enhancer and promoter list was then divided into three equal subgroups labeled as “high”, “medium”, and “low” Pol II-S5P.

### RNA-seq

Quality control and trimming was done as previously described^21^. The reads were mapped with Star (v2.7.3a), then the count was calculated with featureCounts (v2.0.0) at gene level. The differential analysis was done by DESeq2 (v1.26.0) with the software Sartools (v1.7.2)

## Acknowledgements

This work was supported by grants from the INCA (2017-1-PL BIO-02-CEA-1), ANR-23-CE12-0036 and Fondation ARC pour la recherche sur le cancer www.fondation-arc.org.

